# Impaired alanine transport or exposure to D-cycloserine increases the susceptibility of MRSA to β-lactam antibiotics

**DOI:** 10.1101/616920

**Authors:** Laura A. Gallagher, Rebecca K. Shears, Claire Fingleton, Laura Alvarez, Elaine M. Waters, Jenny Clarke, Laura Bricio-Moreno, Christopher Campbell, Akhilesh K. Yadav, Fareha Razvi, Eoghan O’Neill, Alex J. O’Neill, Felipe Cava, Paul D. Fey, Aras Kadioglu, James P. O’Gara

**Affiliations:** School of Natural Sciences, National University of Ireland, Galway, Ireland; Department of Clinical Infection, Microbiology and Immunology, Institute of Infection and Global Health, University of Liverpool, UK; MIMS-Molecular Infection Medicine Sweden, Molecular Biology Department, Umeå University, Umeå, Sweden; Department of Pathology and Microbiology, University of Nebraska Medical Center, Omaha, Nebraska, USA; Department of Clinical Microbiology, Royal College of Surgeons in Ireland, Connolly Hospital, Dublin 15, Ireland; Antimicrobial Research Centre, School of Molecular and Cellular Biology, Faculty of Biological Sciences, University of Leeds, Leeds, UK

**Author notes:** Correspondence: Prof James P. O’Gara, Discipline of Microbiology, School of Natural Sciences, National University of Ireland, Galway, Ireland.

## Abstract

Prolonging the clinical effectiveness of β-lactams, which remain first-line antibiotics for many infections, is an important part of efforts to address antimicrobial resistance. We report here that inactivation of the predicted D-cycloserine (DCS) transporter gene *cycA* re-sensitized MRSA to β-lactam antibiotics. The *cycA* mutation also resulted in hyper-susceptibility to DCS, an alanine analogue antibiotic that inhibits alanine racemase and D-alanine ligase required for D-alanine incorporation into cell wall peptidoglycan (PG). Alanine transport was impaired in the *cycA* mutant and this correlated with increased susceptibility to oxacillin and DCS. The *cycA* mutation or exposure to DCS were both associated with the accumulation of muropeptides with tripeptide stems lacking the terminal D-ala-D-ala and reduced PG crosslinking, prompting us to investigate synergism between β-lactams and DCS. DCS re-sensitised MRSA to β-lactams *in vitro* and significantly enhanced MRSA eradication by oxacillin in a mouse bacteraemia model. These findings reveal alanine transport as a new therapeutic target to enhance the susceptibility of MRSA to β-lactam antibiotics.

## Introduction

Whilst many bacteria can exhibit resistance to select antimicrobials, isolates of the human pathogen *Staphylococcus aureus* can express resistance to all licensed anti-staphylococcal drugs. This results in significant morbidity and mortality, with up to 20% of patients with systemic methicillin resistant *S. aureus* (MRSA) infections dying, despite receiving treatment with anti-staphylococcal drugs [1]. As part of our efforts to identify improved therapeutic approaches for MRSA infections, we recently described the novel use of *β*-lactam antibiotics to attenuate the virulence of MRSA-induced invasive pneumonia and sepsis [2]. We demonstrated that oxacillin-induced repression of the Agr quorum-sensing system and altered cell wall architecture resulted in downregulated toxin production and increased MRSA killing by phagocytic cells, respectively [2]. Supporting this *in vitro* data, a randomised controlled trial involving 60 patients showed that the *β*-lactam antibiotic flucloxacillin in combination with vancomycin shortened the duration of MRSA bacteraemia from 3 days to 1.9 days [3, 4].

Because expression of methicillin resistance in *S. aureus* impacts fitness and virulence and is a regulated phenotype, further therapeutic interventions may also be possible. The complexity of the methicillin resistance phenotype is evident among clinical isolates of MRSA, which express either low-level, heterogeneous (HeR) or homogeneous, high-level methicillin resistance (HoR) [5–7]. Exposure of HeR isolates to *β*-lactam antibiotics induces expression of *mecA,* which encodes the alternative penicillin binding protein 2a (PBP2a) and can select for mutations in accessory genes resulting in a HoR phenotype, including mutations that affect the stringent response and c-di-AMP signalling [8–12]. Because accessory genes can influence the expression of methicillin resistance in MRSA, targeting the pathways associated with such genes may identify new ways to increase the susceptibility of MRSA to β-lactams. To pursue this, we performed a forward genetic screen to identify loci that impact the expression of resistance to β-lactam antibiotics in MRSA. Using the Nebraska Transposon Mutant Library, which comprises 1,952 sequence-defined transposon insertion mutants [13], inactivation of a putative amino acid permease gene, *cycA*, was found to reduce resistance to cefoxitin, the β-lactam drug recommended by the Clinical and Laboratory Standards Institute for measuring *mecA*-mediated methicillin resistance in MRSA isolates. Amino acid transport and susceptibility to oxacillin and D-cycloserine (DCS) were compared in the wild-type and *cycA* mutant grown in chemically defined media (CDM), CDM supplemented with glucose (CDMG) and other complex media. The activity of DCS and *β*-lactams, alone and in combination, against MRSA was measured *in vitro* and in a mouse model of bacteraemia. Peptidoglycan analysis was performed to compare the impact of the *cycA* mutation or exposure to DCS on cell wall structure and crosslinking. Our experiments suggest that therapeutic strategies targeting alanine transport, which was required for resistance to *β*-lactams and DCS, and a re-evaluation of DCS may be important as part of efforts to restore the efficacy of *β*-lactam antibiotics against MRSA.

## Results

### Mutation of *cycA* increases the susceptibility of MRSA to β-lactam antibiotics and D-cycloserine

To identify new ways of controlling expression of methicillin resistance, we sought to identify novel mutations involved in this phenotype. An unbiased screen of the NTML to identify mutants with increased susceptibility to cefoxitin identified NE810 (SAUSA300_1642) (Fig. S1A), which also exhibited a >128-fold increase in susceptibility to oxacillin (Fig. S1B). NE810 was previously identified among several NTML mutants reported to be more susceptible to amoxicillin [14], but was not investigated further. Expression of *mecA* was not affected in NE810 (Fig. S1C) and genome sequence analysis revealed an intact SCC*mec* element and the absence of any other mutations. NE810 was successfully complemented (Fig. S1B), and transduction of the SAUSA300_1642 allele into several MRSA strains from a number of clonal complexes and with different SCC*mec* types was also accompanied by increased cefoxitin and oxacillin susceptibility (Table 1).

**Table 1.** Antibacterial activity (minimum inhibitory concentrations, MIC) and drug synergy (fractional inhibitory concentration indices, ΣFIC) of D-cycloserine (DCS) and several β-lactam antibiotics with different PBP specificity, namely oxacillin (OX; PBP1, 2, 3), nafcillin (NAF; PBP1), cefoxitin (FOX; PBP4), cefaclor (CEC; PBP3), cefotaxime (CTX; PBP2), piperacillin-tazobactin (TZP; PBP3/β-lactamase inhibitor) and imipenem (IMP; PBP1), alone and in β-lactam combinations, against fourteen *S. aureus* strains and *S. epidermidis* RP62A.

| Antibiotic → | OX* | NAF* | DCS* | FOX* | DCS/FOX**<br>(ΣFIC)*** | CEC* | DCS/CEC**<br>(ΣFIC)*** | CTX* | DCS/CTX**<br>(ΣFIC)*** | TZP* | DCS/TZP**<br>(ΣFIC)*** | IPM* | DCS/IPM**<br>(ΣFIC)*** |
| --- | --- | --- | --- | --- | --- | --- | --- | --- | --- | --- | --- | --- | --- |
| ↓ Strain |  |  |  |  |  |  |  |  |  |  |  |  |  |
| JE2 | 64 | 32 | 32 | 64 | 8/8 (0.37) | 64 | 8/2 (0.28) | 64 | 8/4 (0.31) | 32 | 8/0.5 (0.26) | 1 | ND |
| NE810 (*cycA*) | 0.25 | 0.5 | 2 | 8 | ND | 4 | ND | 8 | ND | 2 | ND | 0.125 | ND |
| USA300 | 64 | 32 | 32 | 64 | 8/8 (0.37) | 64 | 8/8 (0.37) | 64 | 8/8 (0.37) | 64 | 8/2 (0.28) | 1 | ND |
| USA300 *cycA* | 0.25 | 1 | 2 | 8 | ND | 4 | ND | 8 | ND | 4 | ND | 0.125 | ND |
| DAR173 | 128 | 128 | 32 | 256 | 4/32 (0.25) | 128 | 8/4 (0.28) | 512 | 8/4 (0.25) | 128 | 4/4 (0.15) | 64 | 4/2 (0.15) |
| DAR173 *cycA* | 0.5 | 8 | 4 | 32 | ND | 16 | ND | 16 | ND | 4 | ND | 1 | ND |
| DAR22 | 128 | 128 | 32 | 128 | 8/8 (0.31) | 128 | 8/8 (0.31) | 512 | 8/4 (0.25) | 128 | 4/4 (0.15) | 128 | 4/1 (0.13) |
| DAR22 *cycA* | 0.5 | 8 | 0.5 | 16 | ND | 16 | ND | 16 | ND | 4 | ND | 0.5 | ND |
| DAR169 | 32 | 32 | 32 | 32 | 8/2 (0.31) | 128 | 4/32 (0.37) | 64 | 4/8 (0.25) | 8 | 4/1 (0.25) | 2 | ND |
| DAR169 *cycA* | 16 | 4 | 0.5 | 16 | ND | 64 | ND | 16 | ND | 2 | ND | 0.5 | ND |
| COL | 512 | 256 | 64 | 512 | 8/128 (0.37) | 128 | 16/32 (0.5) | >2048 | ND | 128 | 32/0.5 (0.5) | 256 | 8/64 (0.37) |
| DAR113 | 128 | 64 | 32 | 128 | 8/8 (0.3) | 64 | 8/4 (0.3) | 256 | 8/8 (0.2) | 128 | 8/4 (0.2) | 16 | 8/0.5 (0.2) |
| BH1CC | 256 | 512 | 32 | 256 | 8/32 (0.3) | 128 | 8/8 (0.3) | 1028 | 8/16 (0.2) | 256 | 8/4 (0.2) | 64 | 8/1 (0.2) |
| BH14B(04) | 128 | 64 | 16 | 128 | 4/32 (0.5) | 128 | 4/32 (0.5) | 512 | 4/32 (0.31) | 64 | 4/1 (0.26) | 64 | 2/8 (0.25) |
| BH8(03) | 128 | 128 | 32 | 128 | 8/32 (0.5) | 256 | 8/64 (0.5) | 128 | 8/16 (0.25) | 128 | 8/8 (0.3) | 64 | 8/0.5 (0.1) |
| BH6 | 128 | 128 | 32 | 128 | 8/16 (0.37) | 256 | 8/64 (0.5) | 512 | 8/32 (0.31) | 128 | 8/4 (0.28) | 32 | 8/0.5 (0.26) |
| DAR202 | 64 | 64 | 32 | 128 | 4/32 (0.37) | 128 | 4/32 (0.37) | 64 | 8/16 (0.5) | 64 | 8/2 (0.28) | 4 | 8/1 (0.5) |
| DAR45 | 2 | 0.5 | 32 | 4 | ND | 32 | 8/2 (0.31) | 4 | ND | 2 | ND | 0.5 | ND |
| DAR13 | 128 | 32 | 32 | 128 | 4/32 (0.37) | 128 | 8/4 (0.28) | 128 | 4/32 (0.37) | 32 | 8/1 (0.28) | 8 | 4/2 (0.37) |
| RP62A | 128 | 2 | 32 | 64 | 6/16 (0.5) | 64 | 6/16 (0.5) | 32 | 8/8 (0.5) | 8 | 8/2 (0.5) | 32 | 8/1 (0.2) |

\* MIC values for each antibiotic when measured individually; μg/ml
\*\* MIC values for each antibiotic when measured in combination, also known as the fractional inhibitory concentration (FIC); μg/ml
\*\*\* FIC indices ( ΣFIC) for antibiotic combinations. ΣFIC = FIC A + FIC B, where FIC A is the MIC of DCS in combination with the β-lactam/MIC of DCS alone, and FIC B is the MIC of the β-lactam in combination with DCS/MIC of the β-lactam alone. Combinations are synergistic when the ΣFIC is $\leq 0.5$ and indifferent when the ΣFIC is $>0.5$ to $<2$ .
<sup>†</sup>ND. Not determined if strain is susceptible (or hyper-resistant) to β-lactam antibiotic or for *cycA* mutants with reduced DCS & β-lactam MICs.

SAUSA300_1642 is annotated as a serine/alanine/glycine transporter with homology to CycA in *Mycobacterium tuberculosis* [15, 16], which influences D-cycloserine (DCS) susceptibility in *Mycobacteria* [15, 16]. In contrast to the observations in *Mycobacteria*, our data showed that NE810 and several unrelated MRSA strains carrying the *cycA* mutation were significantly more susceptible to DCS than the wild type JE2 (Fig. S1D, Table 1). The *cycA* mutation also reduced the DCS MIC of the MSSA strains 8325-4 and ATCC29213 from 32 to 4 μg/ml. DCS inhibits alanine racemase (Alr) that converts L-alanine to D-alanine and the Ddl D-alanyl:D-alanine ligase [17]. A mutant in the putative *ddl* SAUSA300_2039 gene is not available in the NTML library, suggesting that it may be essential. However, the *alr* mutant NE1713 was more susceptible to cefoxitin (Fig. S2A; MIC=16μg/ml) and DCS (Fig. S2B; MIC <0.25μg/ml), consistent with an important role for D-alanine in resistance to both antibiotics.

### CycA is required for alanine transport and D-ala-D-ala incorporation into the peptidoglycan stem peptide

To investigate the role of CycA in amino acid transport, JE2 and NE810 were grown for 8 h in chemically defined media containing 14mM glucose (CDMG) and amino acid consumption in spent media was measured. Although no growth rate or yield difference was noted between JE2 and NE810 in CDMG (Fig. 1A), alanine uptake by NE810 was significantly impaired compared to JE2 (Fig. 1B). Utilisation of other amino acids by NE810 and JE2, including serine and glycine, were similar (Fig. S3). Impaired alanine transport in the *cycA* mutant grown in CDMG correlated with increased susceptibility to oxacillin (1 μg/ml) (Fig. 1C) and DCS (1 μg/ml) (Fig. 1D). These data demonstrate for the first time that CycA in *S. aureus* is required for alanine transport.

**Figure 1.**
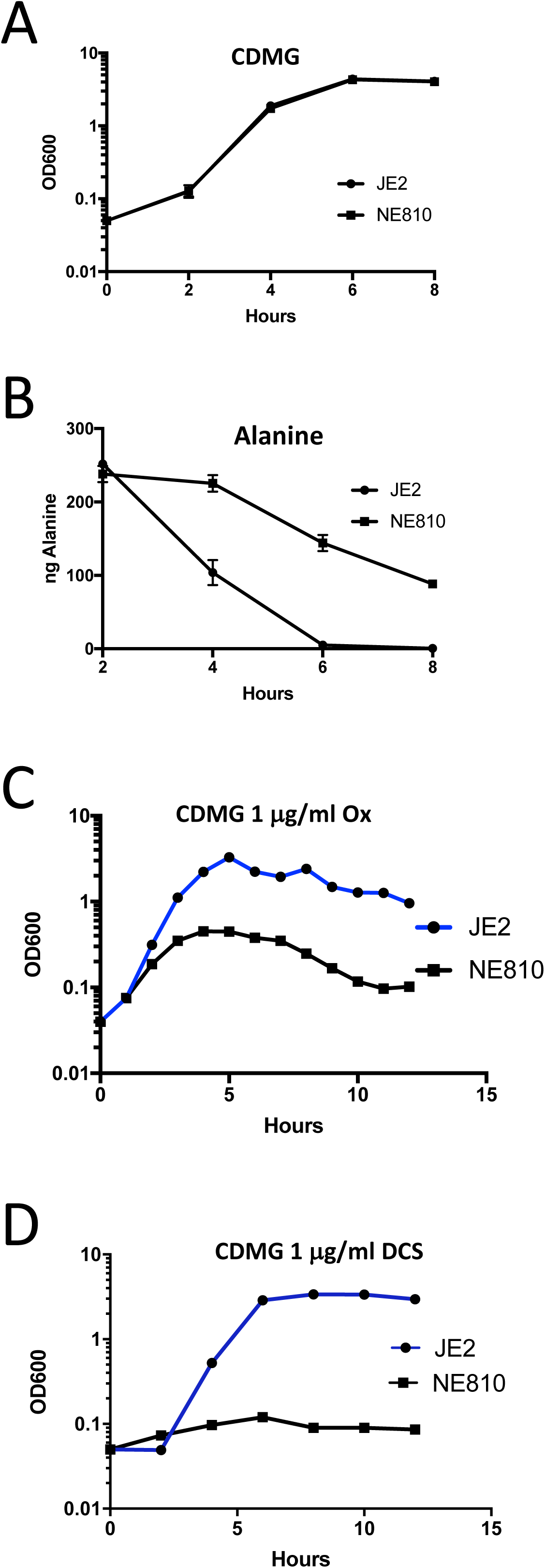
Mutation of *cycA* impairs alanine uptake. **A.** Growth of JE2 and NE810 in chemically defined media supplemented with glucose (CDMG). Cell density was measured at *A*_600_. **B.** Alanine consumption by JE2 and NE810 grown aerobically in CDMG. Residual amino acid was measured in spent media after 2, 4, 6 and 8 h growth. **C and D.** Growth of JE2 and NE810 cultures for 12 h in CDMG supplemented with 1 μg/ml oxacillin (C) or 1 μg/ml D-cycloserine (D). Cell density was measured at *A*_600_.

Quantitative peptidoglycan compositional analysis was performed using UPLC analysis of muramidase-digested muropeptide fragments extracted from exponential phase cultures of JE2 and NE810 grown for 220 mins in TSB media (Fig. S4). The PG profile of the *cycA* mutant revealed a significant accumulation of tripeptides compared to wild-type JE2 (Fig. 2A,B), which was associated with a significant reduction in crosslinking (Fig. 2C). In NE810, the dimer, trimer and tetramer fractions were decreased, which was accompanied by a concomitant increase in the monomer fraction (Fig. 2D). Consistent with this data, exposure of JE2 to DCS 8μg/ml was also associated with a similar accumulation in muropeptides with tripeptide stems (Fig. 2B), reduced cross-linking (Fig. 2C), increased muropeptide monomers and reduced dimers, trimers and tetramers (Fig. 2D). DCS had a strong dose-dependent effect on the accumulation of muropeptides with tripeptide stems, reduced cross-linking and accumulation of monomers (Fig. 2A-D). Sub-inhibitory (0.25*×* MIC) and 4*×* MIC DCS concentrations, were previously shown to be associated with incorporation of an incomplete stem peptide (tripeptide) [17] and reduced D-ala-D-ala levels [18], respectively. These data show that impaired D-ala incorporation in the *cycA* mutant or following exposure to DCS is accompanied by reduced PG cross-linking and increased *β*-lactam susceptibility.

**Figure 2.**
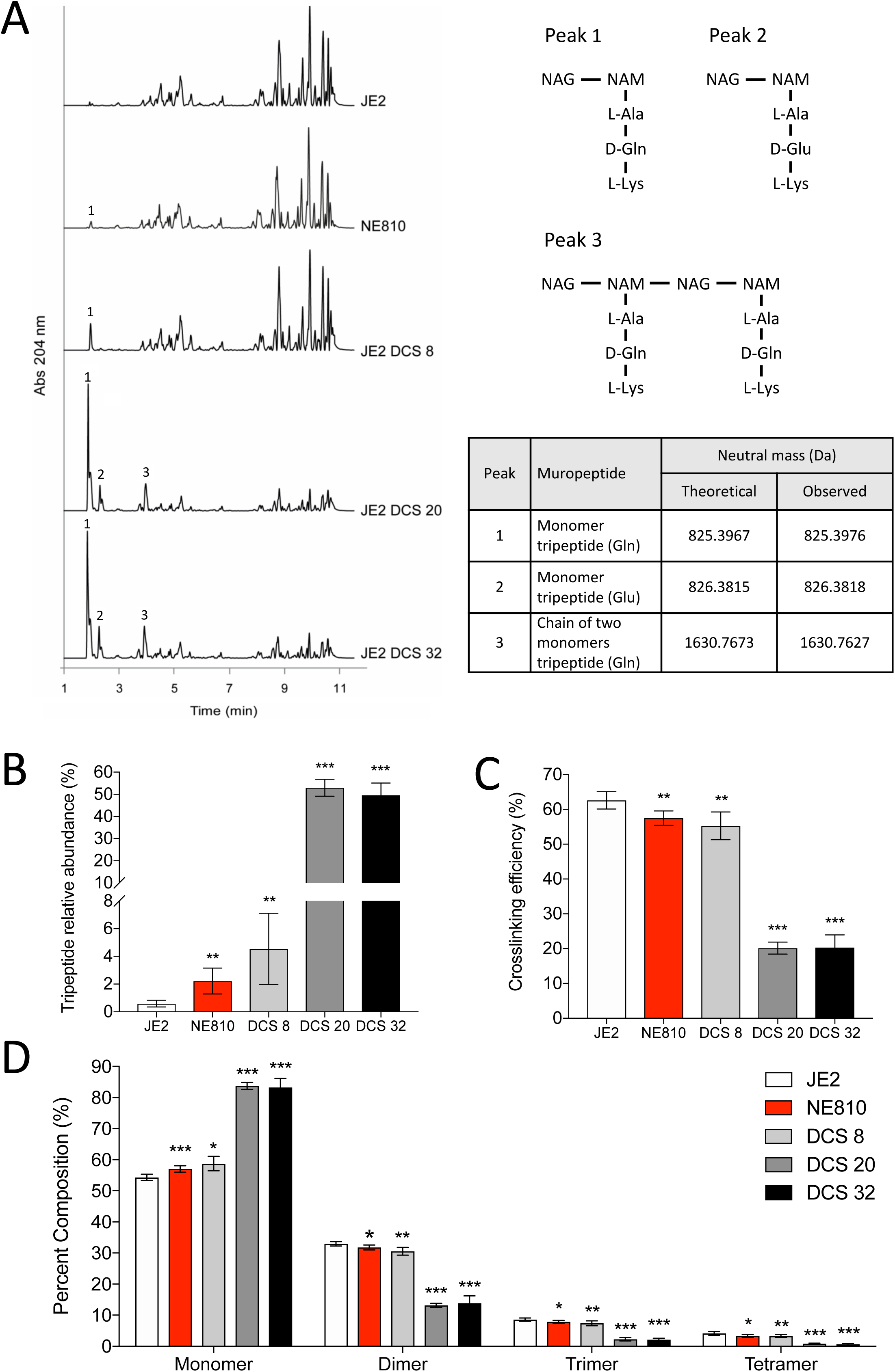
Mutation of *cycA* or D-cycloserine (DCS) treatment impacts peptidoglycan peptide stem length and reduces cell wall crosslinking. **A.** Representative UV chromatograms of peptidoglycan from wild-type JE2, NE810 and JE2 treated with increasing concentrations of DCS (8, 20 and 32 μg/ml). Muropeptides with tripeptide stems are numbered 1-3. The Proposed structures of the three muropeptides with tripeptide stems identified in NE810 and DCS-treated JE2 cells. NAG, N-acetylglucosamine; NAM, N-acetylmuramic acid; L-Ala, L-alanine; D-Gln, D-glutamine; D-Glu, D-glutamic; L-Lys, L-lysine. The theoretical and observed neutral masses determined by MS are indicated. **B.** Relative abundance of muropeptides with tripeptides in the stem. **C.** Relative crosslinking efficiency. **D.** Relative proportions of cell wall muropeptide fractions based on oligomerization. All errors bars represent 95% confidence interval (n = 4). Significant differences determined using Students t-test are denoted using asterisks (* p<0.05; ** p<0.01; *** p<0.001).

### Mutation of *cycA* or exposure to D-cycloserine increases the susceptibility of MRSA to β-lactam antibiotics

Previously reported synergy between DCS and β-lactam antibiotics [17, 19] suggests that impaired alanine uptake in the *cycA* mutant may have the same impact on cell wall biosynthesis as DCS-mediated inhibition of Alr and Ddl. To further investigate this, we compared the activity of DCS and β-lactam antibiotics, alone and in combination, against JE2 and NE810. Checkerboard microdilution assay fractional inhibitory concentration indices (ΣFICs ≤0.5) revealed synergy between DCS and several licensed *β*-lactam antibiotics with different PBP selectivity against JE2 and USA300 FPR3757 (Table 1). Oxacillin and nafcillin were not included in checkerboard assays because measurement of their MICs involves supplementing the media with 2% NaCl, which distorts the MIC of DCS (data not shown). Using the MRSA strains JE2, USA300, DAR173, DAR22, DAR169 and their corresponding *cycA* mutants, the kinetics of killing by DCS, oxacillin and cefoxitin, alone and in combination was measured over 24h using antibiotic concentrations corresponding to 0.125*×*, 0.25*×* and 0.5*×* MICs. Recovery of growth in media supplemented with oxacillin or cefoxitin alone was evident after 8 h (Fig. 3), reflecting the selection and expansion of HoR mutants as described previously [2, 18, 20]. Recovery of growth in cultures exposed to DCS alone was also evident (Fig. 3), which may correlate with our observation that mutants resistant to DCS (on BHI agar supplemented with 128 μg/ml DCS) arise at a rate of approximately 5.5 *×* 10^−8^ per cell per generation. Using combinations of DCS and oxacillin or cefoxitin at 0.125*×* MIC did not achieve a *≥*2 log^10^ reduction in the number of CFU/ml (data not shown). However, at 0.5*×* MIC for strains JE2, USA300, DAR173 and DAR22, DCS (16 μg/ml)/oxacillin (32 μg/ml) and DCS (16 μg/ml)/cefoxitin (32 μg/ml) combinations achieved a *≥*5 log^10^ reduction in the number of CFU/ml compared to oxacillin, cefoxitin or DCS alone (Fig. 3). For strain DAR169, DCS/*β*-lactam combinations at 0.25*×* MIC was sufficient to achieve a *≥*5 log^10^ reduction in CFUs recovered compared to the individual antibiotics (Fig. 3). DCS/*β*-lactam combinations at 0.5*×* MIC were also able to achieve *≥*5 log^10^ reduction in the number of CFU/ml against the methicillin resistant *S. epidermidis* (MRSE) strain RP62A [21] compared to either antibiotic alone (Fig. S5). Checkerboard experiments with fourteen MRSA strains and MRSE strain RP62A further revealed synergy (ΣFICs ≤0.5) between DCS and a range of *β*-lactam antibiotics with different penicillin binding protein (PBP) specificity, namely cefoxitin (PBP4), cefaclor (PBP3), cefotaxime (PBP2), piperacillin-tazobactin (PBP3/*β*-lactamase inhibitor) and imipenem (PBP1) (Table 1).

**Figure 3.**
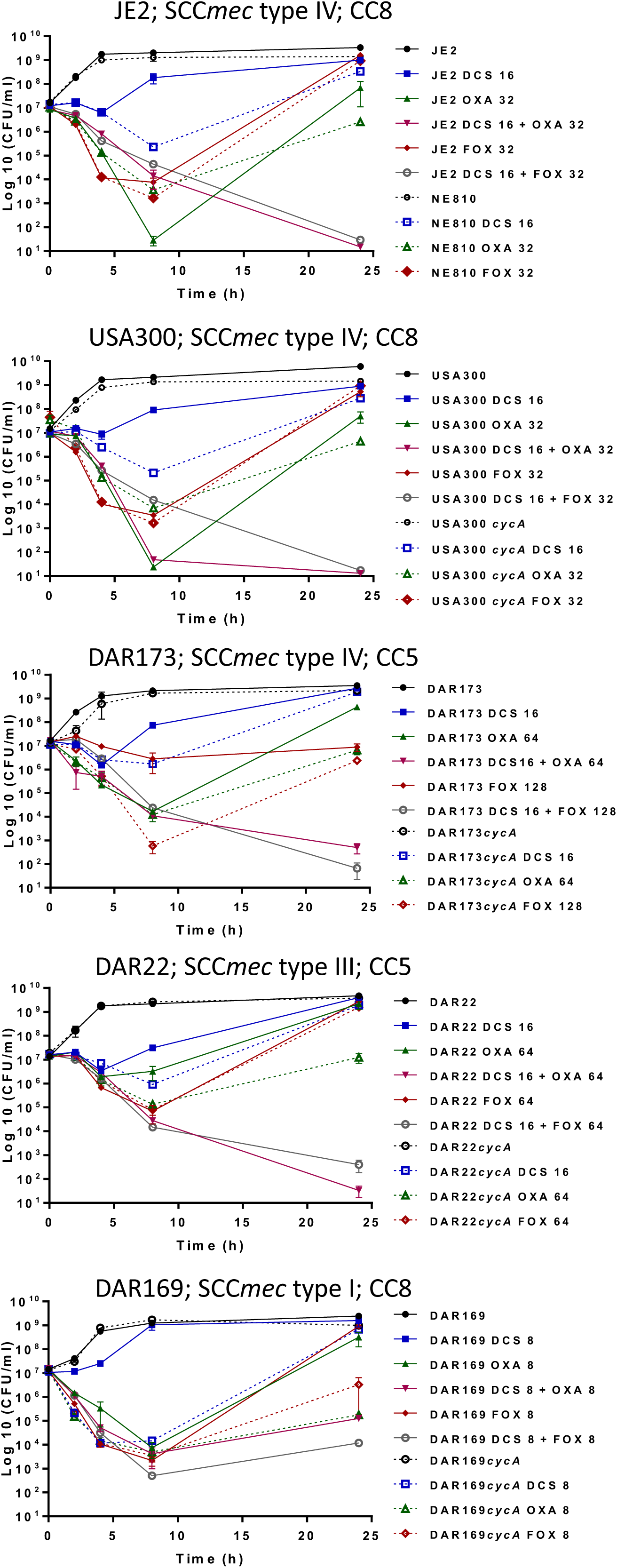
*In vitro* kill curves for D-cycloserine (DCS), oxacillin and cefoxitin with JE2, USA300 FPR3757, DAR173, DAR22, DAR169 and their isogenic *cycA* mutants. Antibiotics at the concentrations indicated (μg/ml) were added to suspensions of overnight bacterial cultures adjusted to 10^7^ CFU/ml in BHI (*A*_600_=0.05), incubated at 37°C and the number of CFU/ml enumerated at 0, 2, 4, 8 and 24 h. The data presented are the mean of three independent experiments, and standard error of the mean is shown. Antibiotic synergism was defined as a ≥2 log^10^ decrease in the number of CFU/ml in cell suspensions exposed to DCS/*β*-lactam combinations compared to the most effective individual antibiotic alone.

This synergy appears to be specific to *β*-lactams and no synergy (ΣFICs > 0.5) was measured between DCS and several antibiotics that are used topically or systemically for the decolonization or treatment of patients colonized/infected with *S. aureus* or MRSA (clindamycin, trimethoprim, mupirocin, ciprofloxacin), or several antibiotics to which *S. aureus* isolates commonly exhibit resistance (tobramycin, kanamycin and spectinomycin) (Table S1). Furthermore the *cycA* mutation had no impact on susceptibility to any of these non-*β*-lactam antibiotics (apart from *ermB*-encoded clindamycin resistance on the transposon).

### Combination therapy with DCS and oxacillin significantly reduces the bacterial burden in the kidneys and spleen of mice infected with MRSA

The virulence of the NE810 mutant and the therapeutic potential of oxacillin in combination with DCS in the treatment of MRSA infections were assessed in mice. Treatment with oxacillin or DCS alone significantly reduced the number of CFUs recovered from the kidneys of mice infected with JE2 (Fig. 4). Furthermore the oxacillin/DCS combination was significantly more effective than either antibiotic alone and the combination was equally effective in reducing the bacterial burden in the kidneys of animals infected with JE2 or NE810 when compared to no treatment (p≤0.0001) (Fig. 4) demonstrating the capacity of DCS to significantly potentiate the activity of *β*-lactam antibiotics against MRSA under *in vivo* conditions. Unexpectedly, oxacillin- or DCS-mediated eradication of NE810 infections in the kidneys was similar to JE2 (Fig. 4). In the spleen, only oxacillin/DCS combination treatment was associated with a significant reduction in the number of CFUs recovered from mice infected with JE2 or NE810 (Fig. S6).

**Figure 4.**
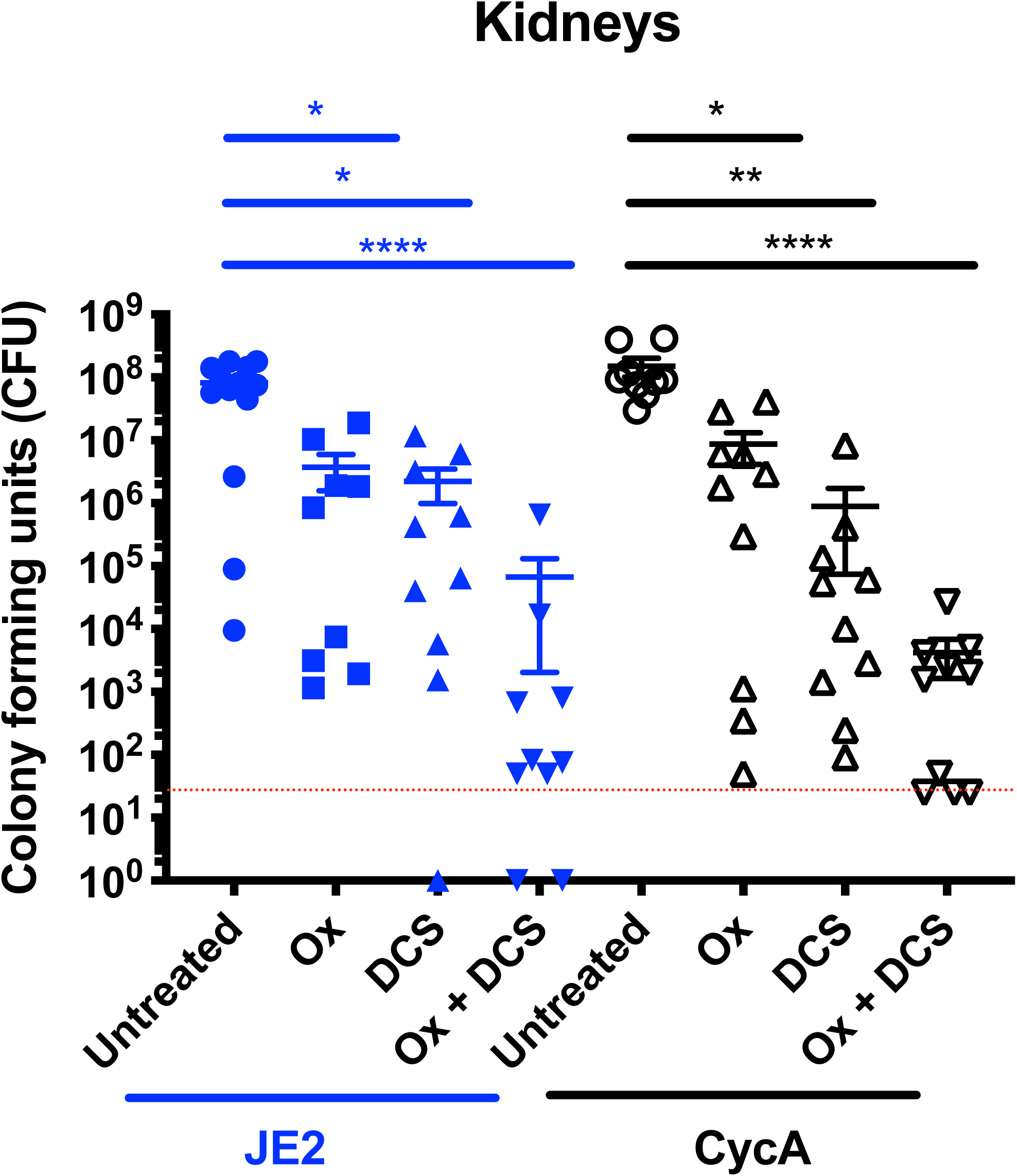
Combination therapy with D-cycloserine and oxacillin significantly reduces the bacterial burden in the kidneys of mice infected with MRSA. The number of colony-forming units (CFU) recovered from the kidneys of mice infected by tail vein injection with 5 × 10^6^ JE2 or NE810 (CycA) and left untreated or treated with 75mg of oxacillin (Ox)/kg, 30mg of DCS/kg or a combination of both Ox and DCS delivered subcutaneously every 12 hours for 5 days. The first antibiotic dose was given 16 hours after infection. Significant differences determined using one-way ANOVA with Kruskal-Wallis test followed by Dunn’s multiple comparisons test are denoted using asterisks (*p≤0.05, **p≤0.01, ****p≤0.0001). The limit of detection (50 colonies) is indicated with a hashed red line.

### Alanine transport and resistance to oxacillin and DCS in chemically defined medium are not dependent on *cycA*

The failure of oxacillin or DCS treatment to enhance the eradication of NE810 infections in the mouse bacteraemia model prompted us to further characterise the growth conditions used for the *in vitro* antibiotic susceptibility assays. Specifically we investigated the role of glucose, which we previously reported to increase the growth requirement for amino acids [22], and which we reasoned may be important for CycA-dependent alanine transport. Growth of JE2 and NE810 was similar in CDM lacking glucose (Fig. 5A) and uptake of alanine (Fig. 5B) and other amino acids (Fig. S7) was unchanged in NE810 compared to JE2. Furthermore NE810 and JE2 grew equally well in CDM supplemented with oxacillin and DSC (Fig. 5C and D). These data explain in part why the *cycA* mutant does not exhibit increased *β*-lactam and DCS susceptibility in the mouse bacteraemia model and further reveal the strong correlation between alanine transport and susceptibility to oxacillin and DCS.

**Figure 5.**
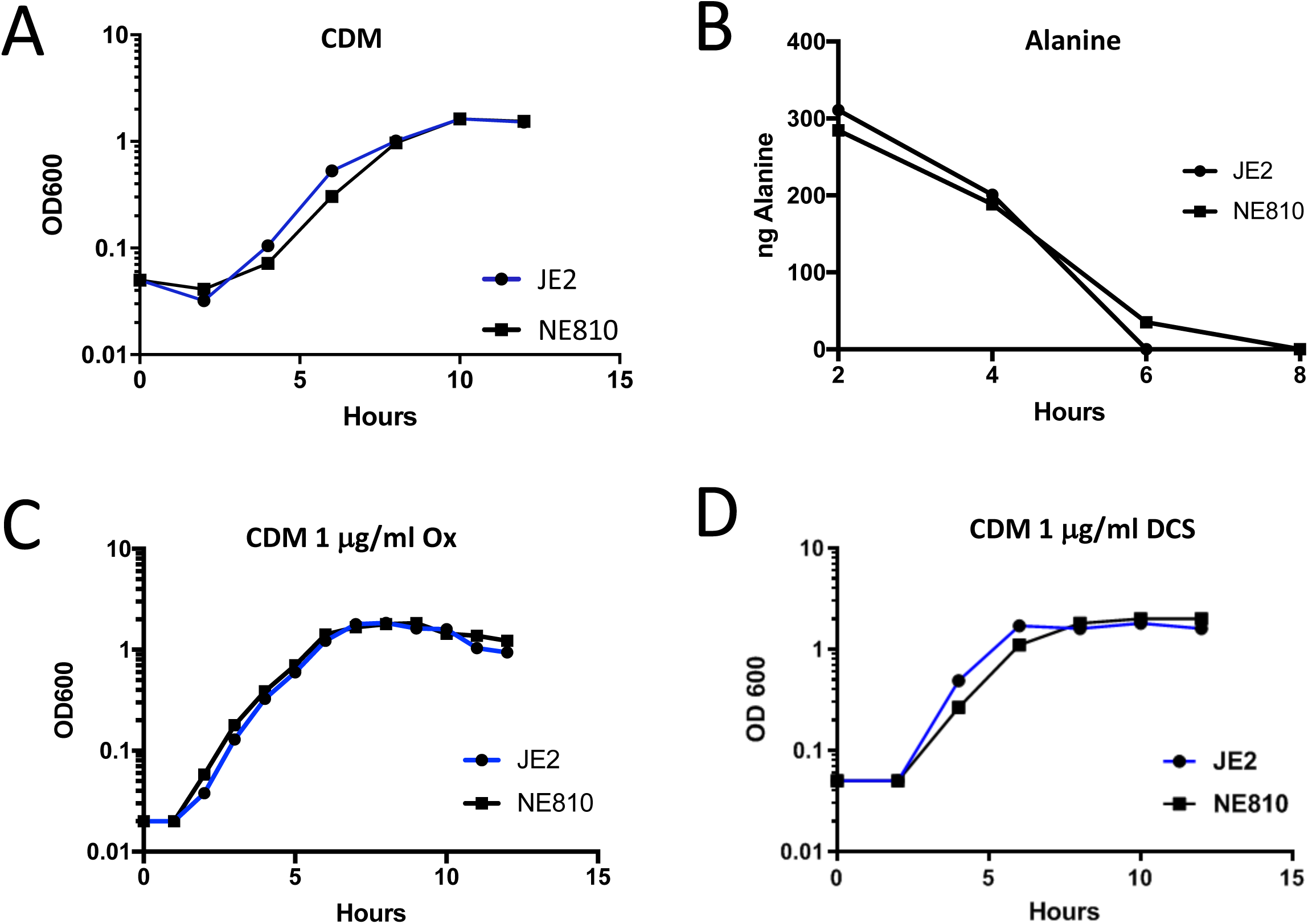
Alanine transport and resistance to oxacillin and D-cycloserine in chemically defined medium are *cycA*-independent. **A.** Growth of JE2 and NE810 in chemically defined medium lacking glucose (CDM). Cell density was measured at *A*_600_. **B.** Alanine consumption by JE2 and NE810 grown aerobically in CDM. Residual amino acid was measured in spent media after 2, 4, 6 and 8 h growth. **C and D.** Growth (cell density at *A*_600_) of JE2 and NE810 cultures for 12 h in CDM supplemented with 1 μg/ml oxacillin (C) or 1 μg/ml DCS (D).

## Discussion

The exploitation of antibiotic re-purposing as part of concerted efforts to address the antimicrobial resistance crisis has been hampered by a lack of mechanistic data to explain demonstrated therapeutic potential and the perception that studies attempting to identify new uses for existing drugs are not hypothesis-driven. In this study, we revealed that CycA was required for full expression of resistance to *β*-lactam antibiotics and DCS. Loss of function of this putative alanine transporter significantly increased the susceptibility of MRSA to *β*-lactam antibiotics, an outcome that could be reproduced through exposure to DCS, which targets the Alr and Ddl enzymes in the early steps of cell wall biosynthesis.

The potential of *β*-lactam/DCS combinations for treatment of MRSA infections follows a recent report that DCS can also potentiate the activity of vancomycin against a laboratory-generated vancomycin highly-resistant *S. aureus* (VRSA) strain *in vitro* and in a silkworm infection model [23]. The excellent safety profile of β-lactam antibiotics makes these drugs particularly attractive as components of combination antimicrobial therapies. When used in the treatment of tuberculosis DCS (trade name Seromycin, The Chao Centre) is typically administered orally in 250 mg tablets twice daily for up to two years. At this dosage, the DCS concentration in blood serum is generally 25-30 μg/ml, which is similar to the concentrations used in our *in vitro* and *in vivo* experiments. The known neurological side effects associated with DCS therapy [24, 25] mean that this antibiotic is unlikely to be considered for the treatment of MRSA infections unless alternative therapeutic approaches have been exhausted. Oxacillin/DCS combination therapy was significantly more effective than DCS or oxacillin alone over a 5-day therapeutic window suggesting that further studies on using DCS to augment *β*-lactams as a treatment option for recalcitrant staphylococcal infections are merited.

Mutation of *cycA* increases the susceptibility of MRSA to *β*-lactam antibiotics and results in hyper-susceptibility to D-cycloserine, whereas a *cycA* point mutation in *M. bovis* contributes, in part, to increased DCS resistance presumably by interfering with transport into the cell [16]. In *E. coli*, *cycA* mutations can also result in increased resistance or have no effect on DCS susceptibility depending on the growth media [26–30], suggesting that CycA is primarily important for DCS resistance under conditions when its contribution to amino acid transport is also important. Our data showing that mutation of *cycA* was not associated with increased DCS resistance strongly suggests that CycA has no role in uptake of this antibiotic in *S. aureus*. Under growth conditions where CycA is required for alanine transport (in nutrient/glucose-replete media), mutation of *cycA* or DCS-exposure have similar effects on the structure of *S. aureus* peptidoglycan (Fig. 8). Consistent with previous studies in *S. aureus* [17] and in *M. tuberculosis* [31], our studies showed a dose-dependent accumulation of muropeptides with a tripeptide stem in MRSA exposed to DCS. The *cycA* mutation was also associated with the increased accumulation of muropeptides with a tripeptide stem. These data indicate that a reduced intracellular alanine pool or inhibition of Alr and Ddl is associated with reduced D-ala-D-ala incorporation into the PG stem peptide. The increased accumulation of tripeptides in turn interferes with normal PBP transpeptidase activity and offers a plausible explanation for increased susceptibility to *β*-lactam antibiotics. The importance of the terminal stem peptide D-ala-D-ala for *β*-lactam resistance has previously been reported. Mutation of the *murF*-encoded ligase, which catalyses of the D-ala-D-ala into the stem peptide also increased *β*-lactam (but not DCS) susceptibility [32, 33]. Similarly growth of a HoR MRSA strain in media supplemented with high concentrations of glycine was accompanied by replacement of the D-ala-D-ala with D-ala-gly and decreased methicillin resistance [34].

Impaired uptake of alanine in CDMG correlated with increased susceptibility to oxacillin and DCS, suggesting that alanine utilisation via CycA is important to make D-alanine available for cell wall biosynthesis and consequently resistance to *β*-lactams. Consistent with this, NE810 also exhibited increased oxacillin susceptibility in BHI, TSB and MH media. However no change in alanine transport or susceptibility to oxacillin and DCS was measured in CDM lacking glucose, which may explain the failure of oxacillin and DCS to more efficiently eradicate NE810 infections in the mouse bacteraemia model. The availability of nutrients such as glucose and amino acids varies in different niches colonised by *S. aureus* during infection ranging from glucose-rich in organs such as the liver [35], to glucose-depleted in established abscesses [36]. In turn this impacts the role of amino acids as carbon sources [22, 37], and potentially the activity of CycA in alanine transport and *β*-lactam susceptibility. Furthermore, normal alanine transport in the *cycA* mutant grown in CDM indicates that an alternative alanine transport mechanism(s) may be active under these growth conditions (Fig. 6). Identification of this alternative alanine permease may be important in the development of therapeutic strategies targeting alanine transport to increase *β*-lactam susceptibility in MRSA, while elucidation of the role of glucose in the control of alanine transport should provide new insights into *β*-lactam resistance.

**Figure 6.**
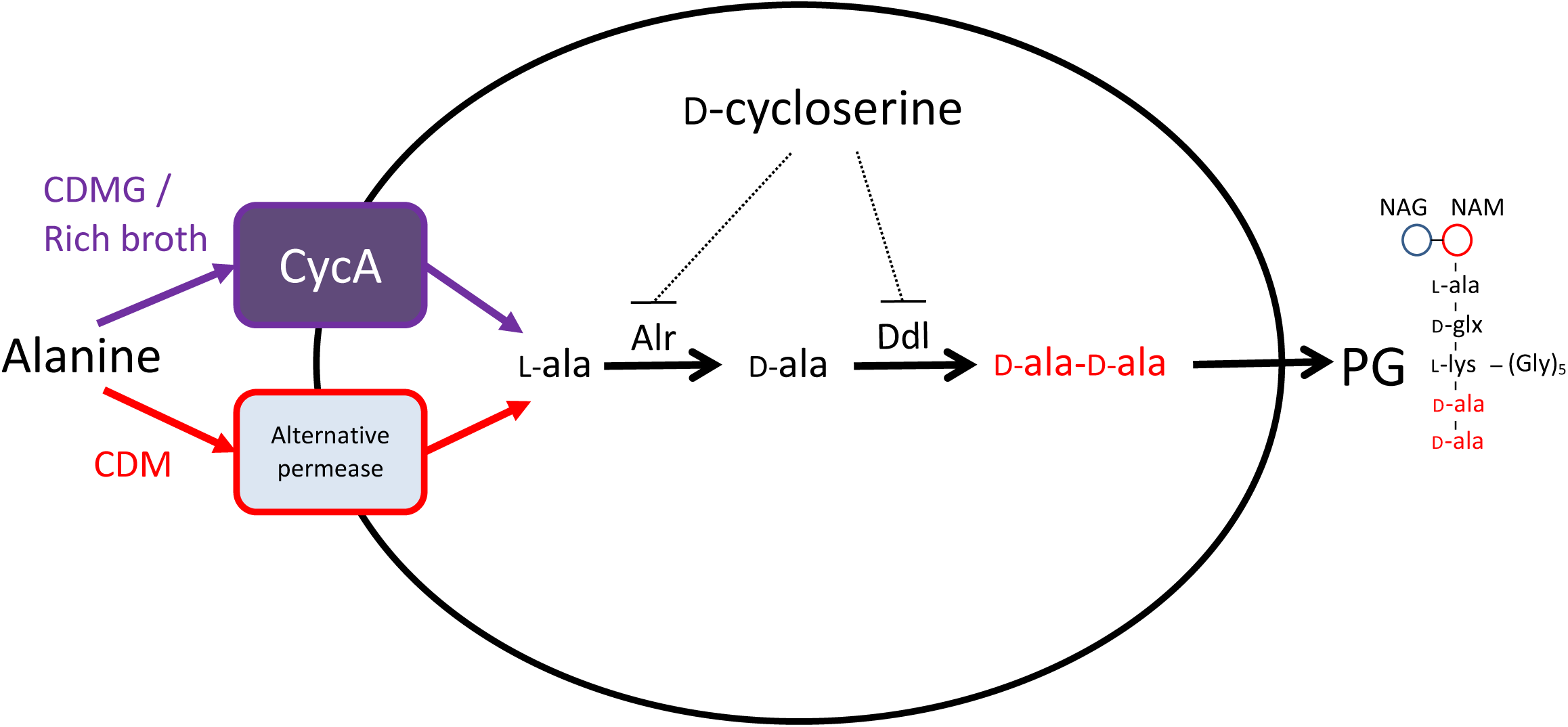
Proposed model depicting how impaired alanine transport associated with mutation of CycA or exposure to DCS can inhibit the D-alanine pathway for peptidoglycan biosynthesis leading to increased susceptibility to *β*-lactam antibiotics.

## Materials and Methods

### Bacterial strains, growth conditions and antimicrobial susceptibility testing

Bacterial strains (Table S2) were grown in Luria Bertoni (LB), brain heart infusion (BHI), Mueller Hinton (MH), nutrient, sheep blood BHI, chemically defined media (CDM) [38] or CDM 14mM glucose (CDMG) [38].

Minimum inhibitory concentrations (MICs) were determined in accordance with CLSI guidelines using plate and broth dilution assays in MH, or MH 2% NaCl for oxacillin and nafcillin. Oxacillin MICs were also measured using E-tests (Oxoid) on MH 2% NaCl. Quality control strains ATCC29213 and ATCC25923 were used for oxacillin and cefoxitin MIC assays, respectively.

### Identification of cefoxitin susceptible MRSA mutant NE810

Cefoxitin (30μg) disks (Oxoid) were used to measure susceptibility of NTML mutants. The zone diameter for JE2 was 18mm NE1868 (*mecA*::Em^r^) was >35mm and NE810 was 22mm. The *cycA* transposon insertion in NE810 was verified by PCR using the primers NE810_Fwd and NE810_Rev (Table S3). Phage 80*α* was used to transduce the NE810 *cycA* allele into JE2 and other strains. Genome sequencing was performed by MicrobesNG using the USA300_FPR3757 genome as a reference. To complement NE810, *cycA* was amplified from JE2 on a 1608 bp fragment using primers NE810F1_Fwd and NE810F1_Rev (Table S3) and cloned into pLI50 using the Clontech In-fusion kit.

### *mecA* transcription analysis

RT-qPCR was performed on a Roche LightCycler with primers mecA1_Fwd and mecA1_Rev for *mecA* and gyrB_Fwd and gyrB_Rev for *gyrB* (internal standard) (Table S3), as described previously [2]. Data presented are the average of three experiments with standard errors.

### Amino acid transport studies

Amino acid analysis in spent media from cultures grown in CDM or CDMG was performed as described previously [22].

### Analysis of peptidoglycan composition in NE810 and JE2 treated with D-cycloserine

Independent quadruplicate 50ml cultures were grown to *A*_600_=0.5, dosed with DCS at 0, 8, 20 or 32 μg/ml for 100 mins, then harvested and resuspended in 5ml PBS (Fig. S4) before peptidoglycan was extracted as described previously [39]. Mass spectrometry was performed on a Waters XevoG2-XS QTof mass spectrometer. Structural characterization of muropeptides was determined based on their MS data and MS/MS fragmentation pattern, matched with PG composition and structure reported previously [34, 40–42].

### Antibiotic synergy analysis using the microdilution checkerboard assay

Antibiotic synergism was measured using the checkerboard microdilution method in 96-well plates inoculated with 5 × 10^5^ CFU/ml. Growth or no growth was recorded after 24 h at 37°C. The fractional inhibitory concentration index (ΣFIC) was calculated for each drug combination in triplicate experiments with an FIC index ≤0.5 considered synergistic.

### Kill curve assays

Overnight cultures adjusted to 10^7^ CFU/ml were exposed to 0.125*×*, 0.25*×*, and 0.5*×* MIC of oxacillin, cefoxitin and DCS alone or in combination, and the number of colony forming units (CFU)/ml enumerated at 0, 2, 4, 8 and 24 h. Data is presented at the antibiotic concentrations where synergy was measured i.e. 0.5*×* MIC for JE2, USA300, DAR173, DAR22, DAR113, BH1CC, and RP62A, and 0.25*×* MIC for DAR169. Synergism was defined as a ≥2 log^10^ decrease in the number of CFU/ml in cell suspensions exposed to DCS/*β*-lactam combinations compared to the most effective individual drug after 8 h.

### Mouse infection experiments

6-8 week-old, age matched, outbred CD1 female mice (Charles River, UK) were used in a non-lethal model of bacteremia. JE2 and NE810 cultures were grown to *A*_600_=0.5 in BHI, washed in PBS, adjusted to 1×10^8^ CFU/ml. Mice were infected intravenously (via the tail vein) with 5×10^6^ CFU (*n* = 10 mice per group). The infections were left untreated (PBS control) or treated with either 75mg oxacillin/Kg/12 hours, 30mg DCS/Kg/12 hours or a combination of both (first antibiotic dose administered 16 hours post infection), before being sacrificed after 5 days. Bacteria present in homogenised spleens and kidneys recovered from the mice were enumerated on blood agar.

### Ethics Statement

Mouse experiments were approved by the UK Home Office (License Number 40/3602) and the University of Liverpool Animal Welfare and Ethics Committee. This study was carried out in strict accordance with the UK Animals (Scientific Procedures) Act 1986. All efforts were made to minimize suffering.

### Statistical analysis

Two-tailed Student’s t-Tests and one-way ANOVA with Kruskal-Wallis test followed by Dunn’s multiple comparisons test in the GraphPad Prism application (for the mouse infection experiments) were used to determine statistically significant differences in assays performed during this study. A *p* value <0.05 was deemed significant.

## Acknowledgements

We thank Craig Winstanley and Kate Reddington for generously providing bacterial strains.

## Footnotes

### Disclaimer

The funders had no role in study design, data collection and analysis, decision to publish, or preparation of the manuscript.

## SUPPLEMENTARY DATA

### Supplementary Tables

**Table S1.**
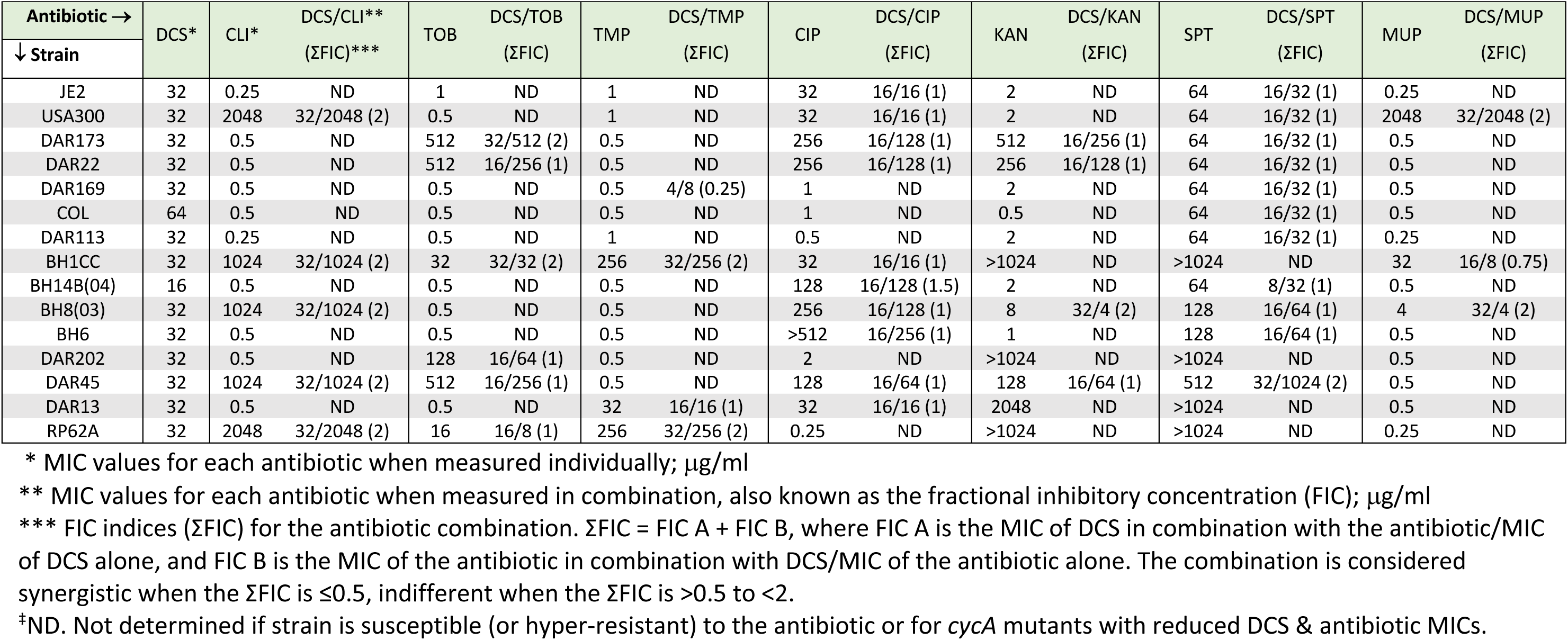
Antibacterial activity (minimum inhibitory concentrations, MIC) and drug synergy (fractional inhibitory concentration indices, ΣFIC) of D-cycloserine (DCS), and clindamycin (CLI), tobramycin (TOB), trimethoprim (TMP), mupirocin (MUP), ciprofloxacin (CIP), kanamycin (KAN) and spectinomycin (SPT) alone and in DCS/antibiotic combinations, against fourteen *S. aureus* strains and *S. epidermidis* RP62A.

**Table S2.** Bacterial strains and plasmids used in this study

| Strains/plasmids | Relevant Details |
| --- | --- |
| RN4220 | Restriction-deficient laboratory *S. aureus* |
| USA300 FPR3757 | Community associated MRSA isolate of the USA300 lineage [43]. SCCmec type IV. CC8. |
| JE2 | USA300 cured of p01 & p03. Parent of Nebraska Transposon Mutant Library (NTML). |
| NE810 | JE2 NTML *cycA* (SAUSA300_1642) mutation. Erm<sup>r</sup>. |
| USA300 *cycA* | USA300 FPR3757 *cycA*. Constructed by transduction of *cycA*::Tn allele from NE810. |
| NE1868 | JE2 NTML *mecA* mutation. |
| NE1713 | JE2 NTML *alr* (SAUSA300_2027) mutation. |
| BH1CC | MRSA clinical isolate; SCCmec type II; CC8 [44] |
| COL | MRSA reference strain; SCCmec type I; CC8 [45] |
| BH14(04) | MRSA clinical isolate; SCCmec type IV; CC22 [44] |
| BH8(03) | MRSA clinical isolate; SCCmec type IV; CC22 [44] |
| BH6(03) | MRSA clinical isolate; SCCmec type II; CC8 [44] |
| DAR113 | MRSA reference isolate; SCCmec type IV; CC22 [44, 46] |
| DAR13 | MRSA reference isolate; SCCmec type IV; CC8 [44, 46] |
| DAR45 | MRSA reference isolate; SCCmec type II; CC30 [44, 46] |
| DAR202 | MRSA reference isolate; SCCmec type III; CC239 [44, 46] |
| DAR173 | MRSA reference isolate; SCCmec type IV; CC5 [44, 46] |
| DAR173 *cycA* | DAR173 *cycA* mutant (*cycA*::Tn allele from NE810) |
| DAR22 | MRSA reference isolate; SCCmec type III; CC5 [44, 46] |
| DAR22 *cycA* | DAR22 *cycA* mutant (*cycA*::Tn allele from NE810) |
| DAR169 | MRSA Reference strain; SCCmec type I; CC8 [44, 46] |
| DAR169 *cycA* | DAR169 *cycA* mutant (*cycA*::Tn allele from NE810) |
| 8325-4 | NCTC 8325 derivative cured of prophages [47], methicillin susceptible, CC8. |
| 8325-4 *cycA* | 8325-4 *cycA* mutant (*cycA*::Tn allele from NE810). |
| ATCC 29213 | Methicillin susceptible *S. aureus* strain for antibiotic susceptibility testing. |
| ATCC 29213 *cycA* | ATCC 29213 *cycA* mutant (*cycA*::Tn allele from NE810). |
| ATCC 25923 | Methicillin susceptible *S. aureus* strain for antibiotic susceptibility testing. |
| *S. epidermidis* RP62A | ATCC 35984. Methicillin resistant, biofilm positive. [21] |
| *E. coli* | *E. coli* HST08 |
| **Plasmids** |  |
| pLI50 | *E. coli-Staphylococcus* shuttle vector. Ap<sup>r</sup> (*E. coli*), Cm<sup>r</sup>. (*Staphylococcus*) |
| p*cycA* | pLI50 carrying *cycA* from JE2 |

**Table S3.**
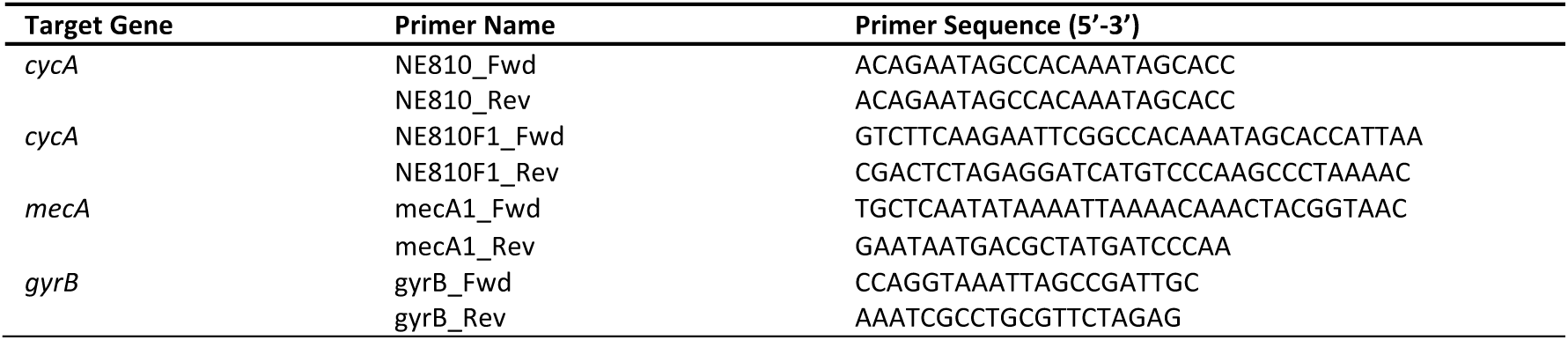
Oligonucleotide primers used in this study

### Supplementary Figures

**Figure S1.**
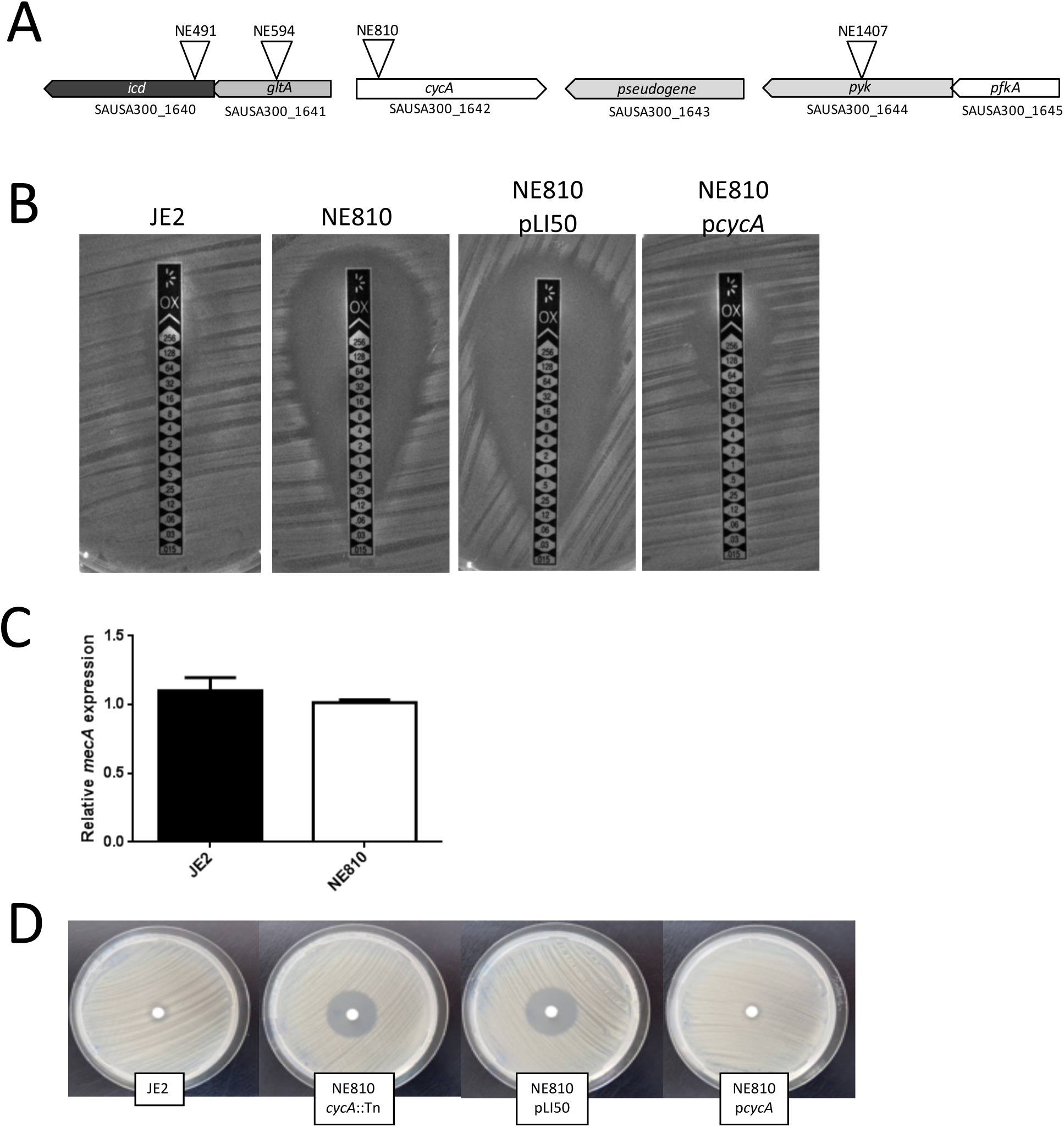
Mutation of *cycA* increases the susceptibility of MRSA to β-lactam antibiotics and D-cycloserine. **A.** Chromosomal location of *cycA* and neighbouring genes *icd* (isocitrate dehydrogenase), *gltA* (citrate synthase), *pyk* (pyruvate kinase) and *pfkA* (6-phosphofructokinase). The locations of transposon insertions in NE810, NE491, NE594 and NE1497 mutants from the Nebraska library are indicated. **B.** E-test measurement of oxacillin minimum inhibitory concentrations (MICs) in JE2 (wild type), NE810 (*cycA*::Tn), NE810 carrying pLI50 (control) and p*cycA*. **C.** Comparison of relative *mecA* gene expression by LightCycler RT-PCR in JE2 and NE810 grown to *A*600=3 in BHI media. The data are the average of three independent experiments and standard deviations are shown **D.** Comparison of zones of inhibition around D-cycloserine 30µg disks on lawns of JE2, NE810 (*cycA*::Tn), NE810 pLI50 (control) and NE810 p*cycA* grown on Mueller Hinton (MH) agar.

**Figure S2.**
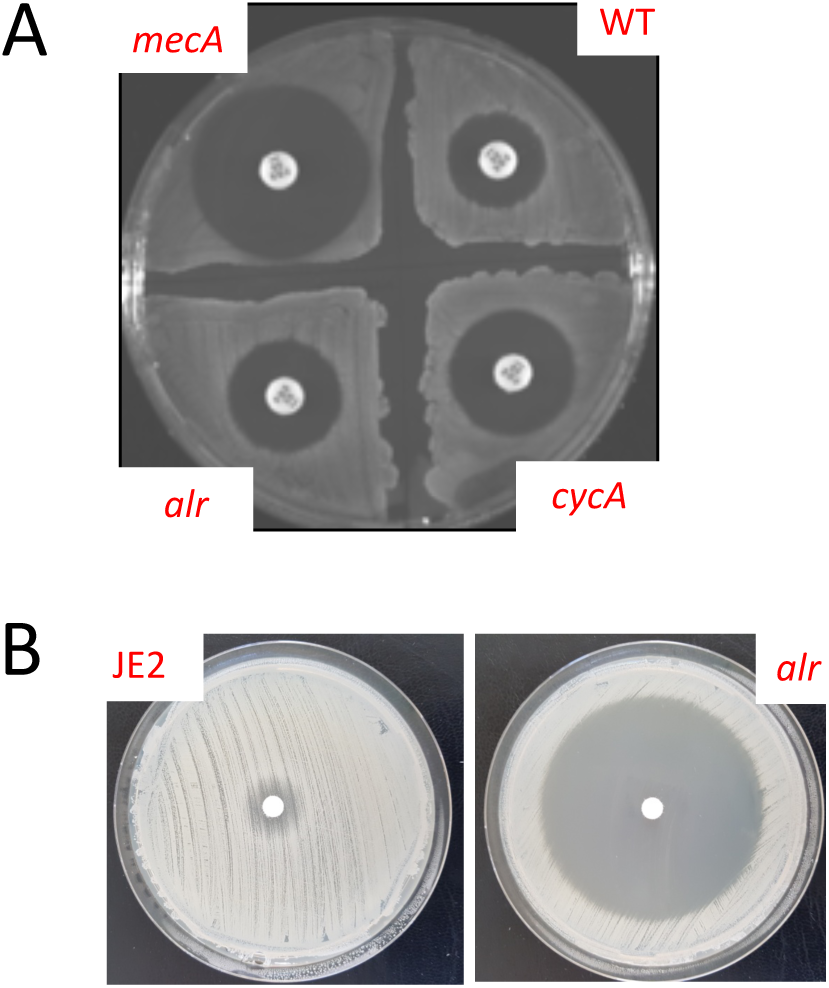
**A.** Susceptibility of JE2 (wild type), NE1868 (*mecA*::Tn), NE810 (*cycA*::Tn), and NE1713 (*alr*::Tn) grown on MH agar to cefoxitin (FOX, 30μg disks). **B.** Susceptibility of JE2 (wild type) and NE1713 (*alr*::Tn) grown on MH agar to D-cycloserine (DCS, 30μg disks).

**Figure S3.**
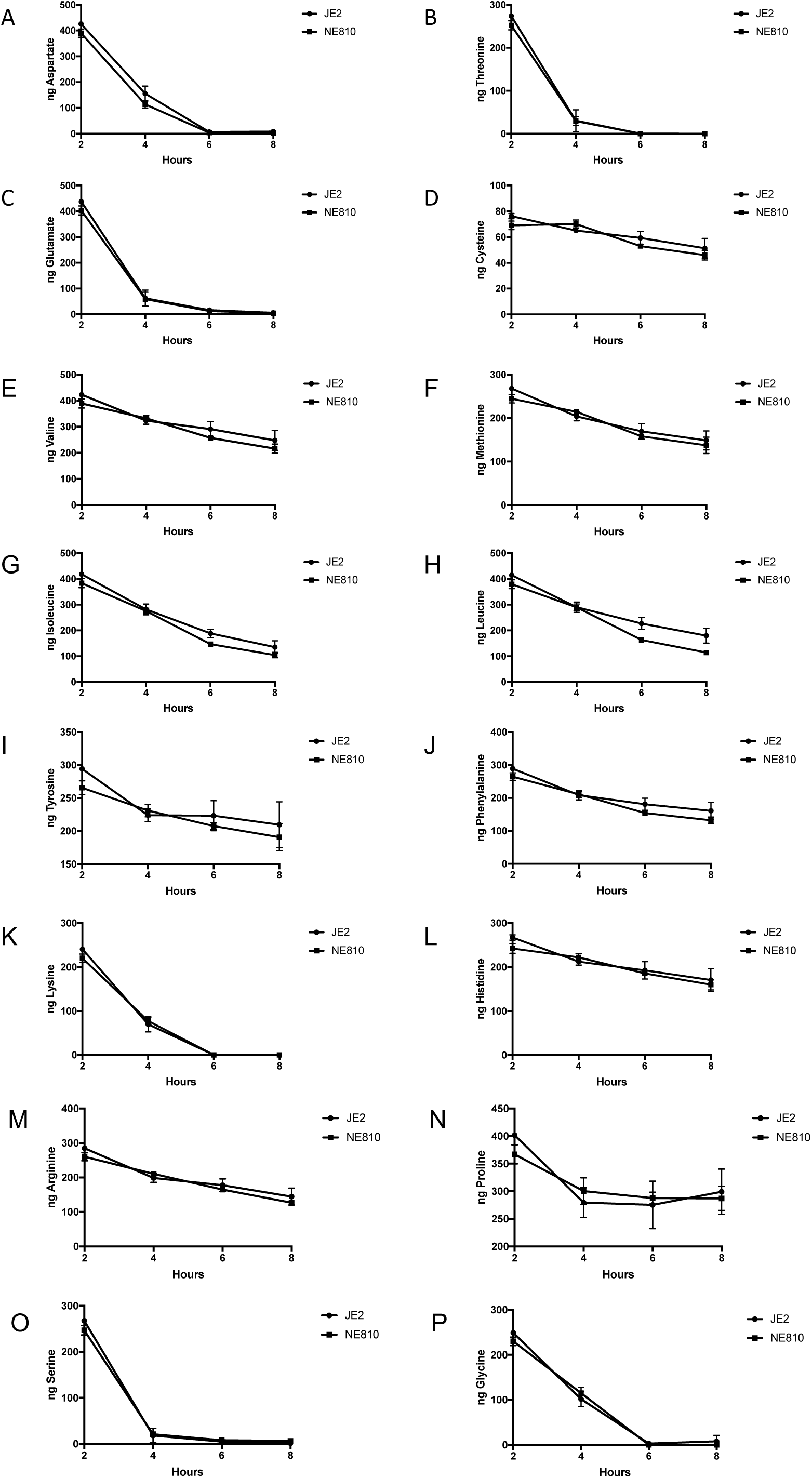
Amino acid consumption by JE2 and NE810 grown aerobically in chemically defined media containing 14mM of glucose (CDMG). Residual amino acids were measured in spent media after 2, 4, 6 and 8 h growth.

**Figure S4.**
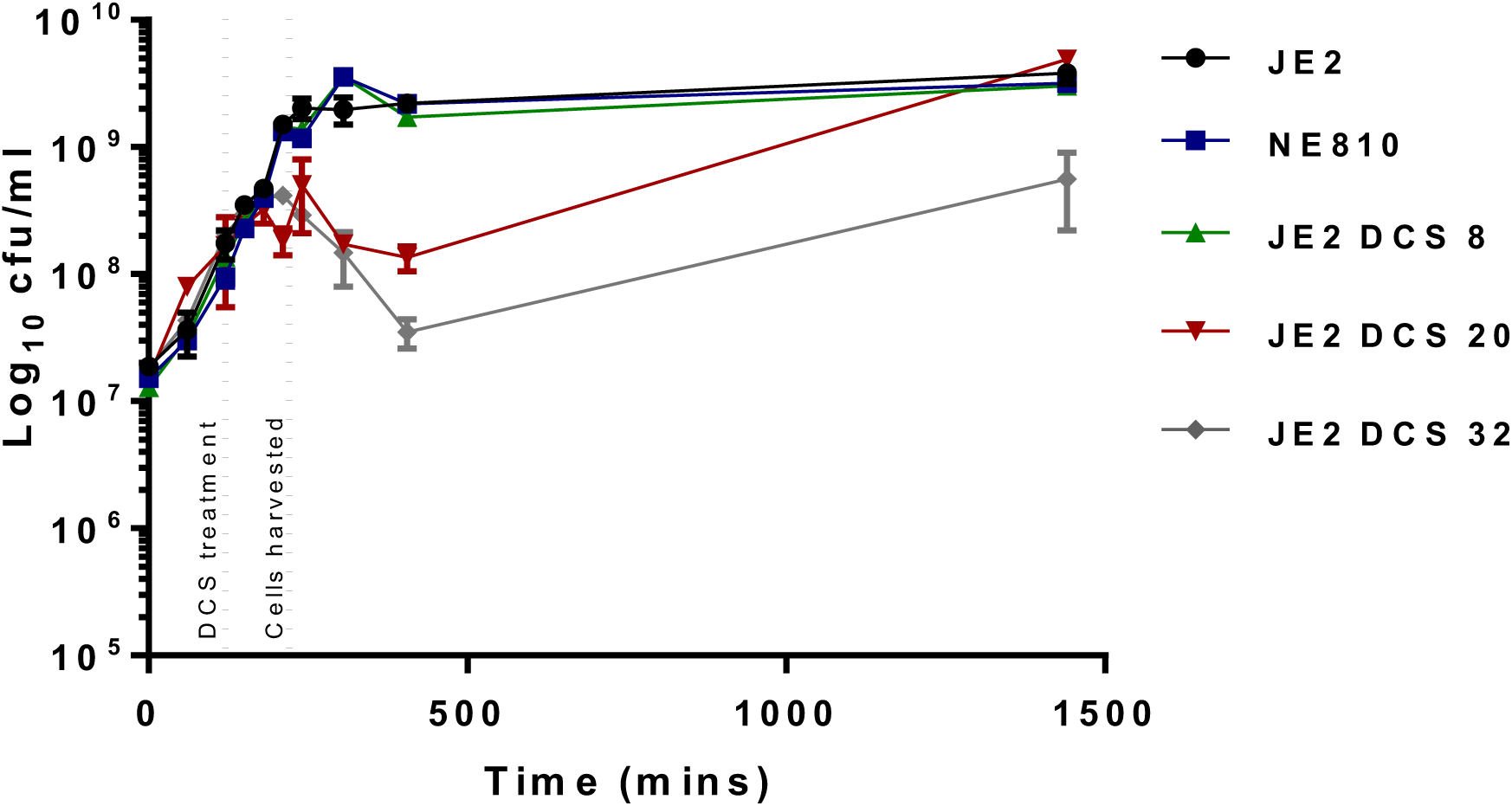
Preparation of cell suspensions for peptidoglycan extraction and structural analysis by UPLC-MS. 50 ml flask cultures were inoculated into fresh BHI media from overnight cultures at a starting cell density of *A*600=0.05 and incubated at 37°C. The number of CFU/ml was enumerated every 1-2 h for 6 h and again after 24 hours. For JE2 cultures being dosed with DCS, the antibiotic was added after approximated 2 h (*A*600≈0.5) and the cells collected after a further 100 mins. Cells from untreated JE2 and NE810 control cultures were collected at the same time point.

**Figure S5.**
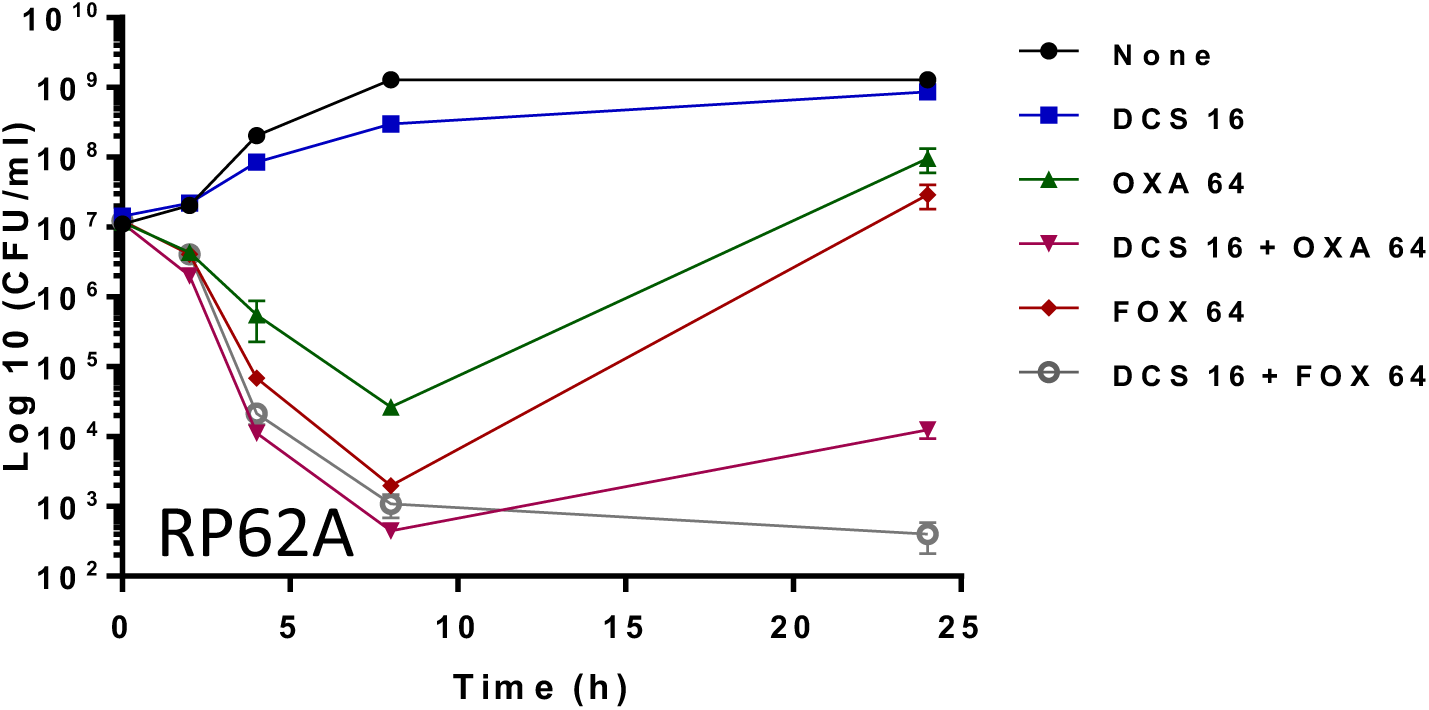
*In vitro* kill curves for D-cycloserine (DCS), oxacillin and cefoxitin with methicillin resistant *S. epidermidis strain* RP62A. Antibiotics at the concentrations indicated (equivalent to 0.5×MIC) were added to suspensions of overnight bacterial cultures adjusted to 10^7^CFU/ml in BHI, incubated at 37°C and the number of CFU/ml enumerated at 0, 2, 4, 8 and 24 h. The data presented are the mean of three independent experiments. Antibiotic synergism was defined as a ≥2 log^10^ decrease in the number of CFU/ml in cell suspensions exposed to DCS/β-lactam combinations compared to the most effective individual antibiotic alone.

**Figure S6.**
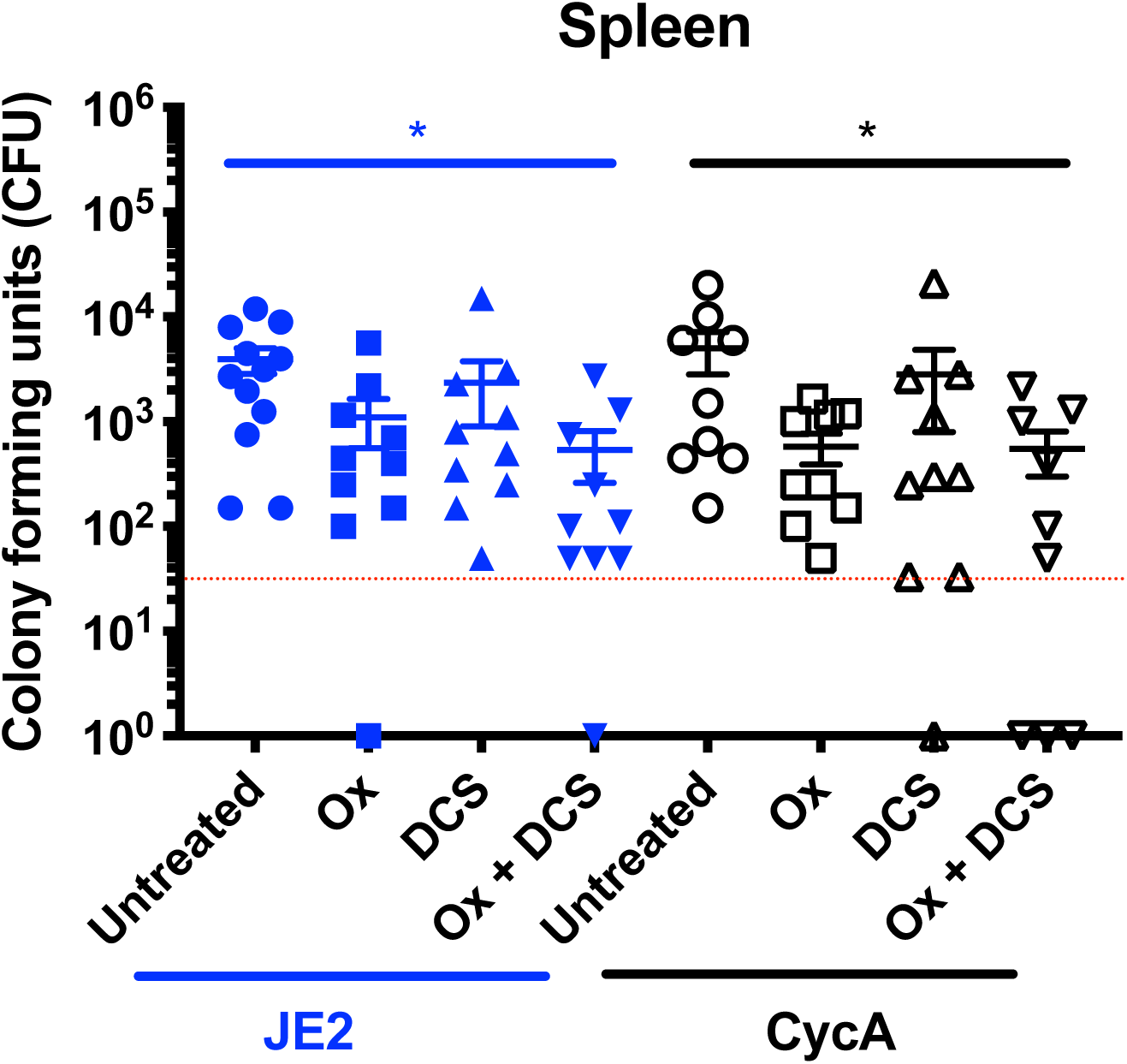
Combination therapy with D-cycloserine and oxacillin significantly reduces the bacterial burden in the spleen of mice infected with MRSA. The number of colony-forming units (CFU) recovered from the spleens of mice infected by tail vein injection with 5 × 10^6^ JE2 or NE810 (CycA) and left untreated or treated with 75mg of oxacillin (Ox)/kg, 30mg of DCS/kg or a combination of both Ox and DCS delivered subcutaneously every 12 hours for 5 days. The first antibiotic dose was given 16 hours after infection. Significant differences determined using one-way ANOVA with Kruskal-Wallis test followed by Dunn’s multiple comparisons test are denoted using asterisks (*p≤0.05). The limit of detection (50 colonies) is indicated with a hashed red line.

**Figure S7.**
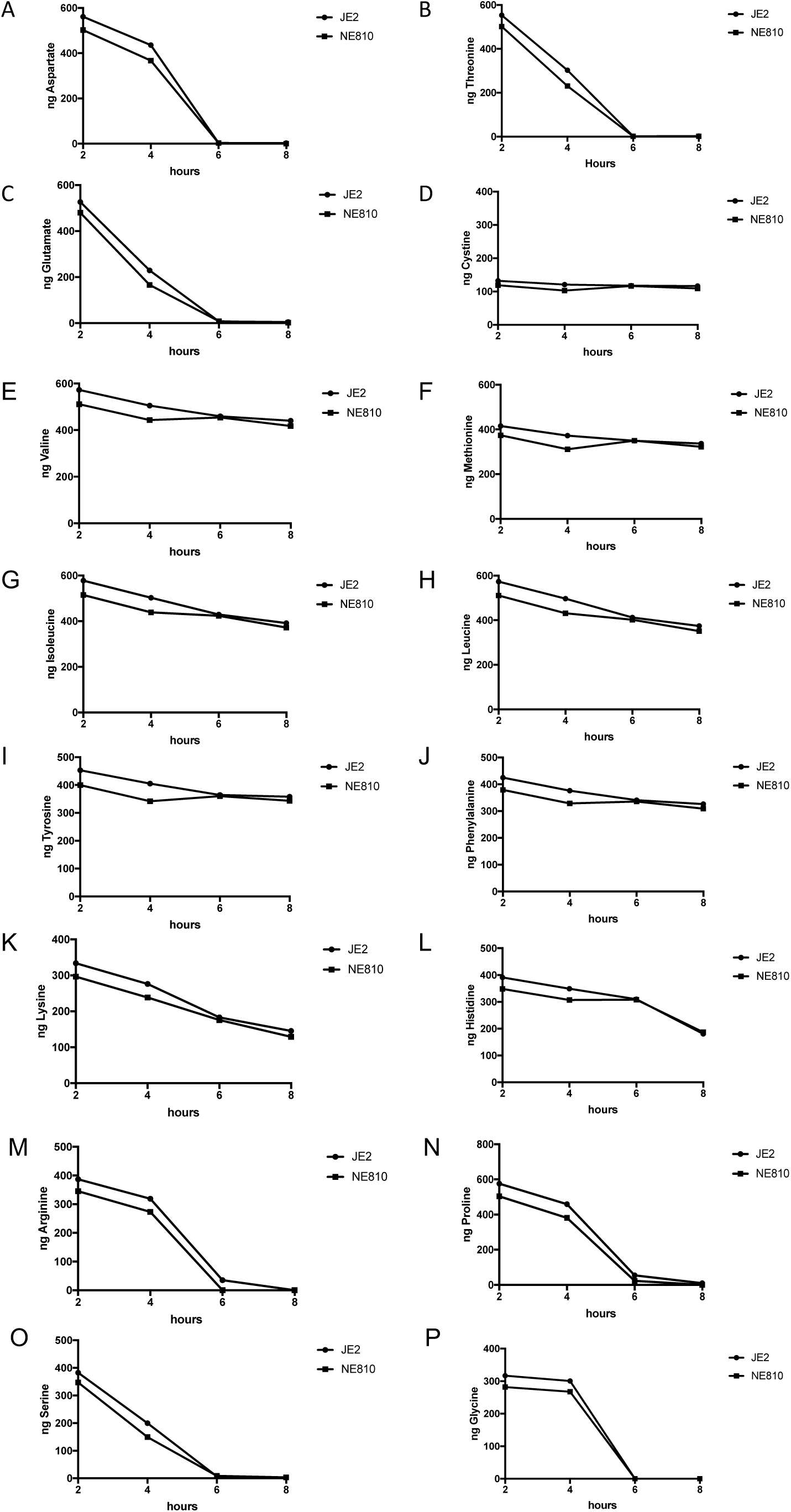
Amino acid consumption by JE2 and NE810 grown aerobically in chemically defined media lacking glucose (CDM). Residual amino acids were measured in spent media after 2, 4, 6 and 8 h growth.

